# Pre-clinical evaluation of antiviral activity of nitazoxanide against Sars-CoV-2

**DOI:** 10.1101/2021.12.17.473113

**Authors:** Jean-Sélim Driouich, Maxime Cochin, Franck Touret, Paul-Rémi Petit, Magali Gilles, Grégory Moureau, Karine Barthélémy, Caroline Laprie, Thanaporn Wattanakul, Palang Chotsiri, Richard M. Hoglund, Joel Tarning, Fanny Escudié, Ivan Scandale, Eric Chatelain, Xavier de Lamballerie, Caroline Solas, Antoine Nougairède

## Abstract

To address the emergence of SARS-CoV-2, multiple clinical trials in humans were rapidly started, including those involving an oral treatment by nitazoxanide, despite no or limited pre-clinical evidence of antiviral efficacy. In this work, we present a complete pre-clinical evaluation of the antiviral activity of nitazoxanide against SARS-CoV-2. First, we confirmed the *in vitro* efficacy of nitazoxanide and tizoxanide (its active metabolite) against SARS-CoV-2. Then, we demonstrated nitazoxanide activity in a reconstructed bronchial human airway epithelium model. In a SARS-CoV-2 virus challenge model in hamsters, oral and intranasal treatment with nitazoxanide failed to impair viral replication in commonly affected organs. We hypothesized that this could be due to insufficient diffusion of the drug into organs of interest. Indeed, our pharmacokinetic study confirmed that concentrations of tizoxanide in organs of interest were always below the *in vitro* EC_50_. These preclinical results suggest, if directly applicable to humans, that the standard formulation and dosage of nitazoxanide is not effective in providing antiviral therapy for Covid-19.

## Introduction

The threat of a global pandemic caused by a virus from the *Coronaviridae* family, which are enveloped positive-stranded RNA viruses, has been hanging over the whole world since the emergence of the severe acute respiratory syndrome coronavirus (SARS-CoV) and the Middle East respiratory syndrome (MERS-CoV). In December 2019, cases of pneumonia were reported in Wuhan, China, (1). Few months later, the causative agent was identified as a new betacoronavirus (2). Named SARS-CoV-2, this pathogen progressed worldwide to such an extent that its disease, called coronavirus disease 2019 (COVID-19), was characterized as a pandemic by the World Health Organization on March 2020 (3). COVID-19 leads to a broad spectrum of clinical syndromes, ranging from pauci-symptomatic disease to severe pneumonia and acute respiratory distress syndrome (4). To date, there are no approved small molecules targeting coronavirus viral replication.. Therefore, drug repurposing has been considered as an interesting strategy to find an active antiviral therapy against SARS-CoV-2.

Nitazoxanide (NTZ) was originally developed as an antiprotozoal agent and marketed for the treatment of *Giardia* and *Cryptosporidium* infections. In recent years, it was identified as a broad-spectrum antiviral drug (5, 6). NTZ, and its active circulating metabolite, tizoxanide (TIZ), inhibit the replication of a wide range of RNA and DNA viruses in cell culture assays including hepatitis B, hepatitis C, rotavirus, norovirus, dengue, yellow fever, Japanese encephalitis virus and the human immunodeficiency virus (6-8). Its inhibitory activity against viruses inducing respiratory infections was specifically investigated (9). Notably, NTZ possesses *in vitro* antiviral activity against influenza virus by blocking the maturation of the viral hemagglutinin, as well as against MERS coronavirus and other coronaviruses by inhibiting expression of the viral N protein (8, 10-12).

It is thus quite naturally that this molecule was rapidly considered as a potential repurposing candidate for COVID-19 management (13-18). NTZ was one of the first molecules studied *in vitro* against SARS-CoV-2. One of the earliest studies on SARS-CoV-2 reported a 50% effective concentration (EC_50_) of 2.12μM in Vero E6 cells at 48 h post-infection (19). Assumptions regarding the possible role of TIZ against numerous targets involved in SARS-CoV-2 pathogenesis affecting viral entry and multiplication were rapidly proposed (20). Additionally, recent findings have also demonstrated that NTZ could inhibit the TMEM16 protein, a calcium-activated ion channel involved in phospholipid transposition between the cell membranes, and block SARS-CoV-2-Spike induced syncytia (21). In addition, NTZ may have the capacity to boost host innate immune responses, affecting the well-described COVID-19 inflammatory cytokine storm. Ambitious expectations have also been raised about its potential ability to improve multi-organ damage and providing added value to patients with comorbidities (20). Consequently, many clinical trials in human investigating the efficacy and the safety of an oral treatment of NTZ alone or in combination with other anti-SARS-CoV-2 candidates are ongoing worldwide (https://clinicaltrials.gov/; search terms: nitazoxanide | Covid19).

However, a pre-clinical *in vivo* investigation of the activity of NTZ against SARS-CoV-2 had yet to be conducted. In the present study, we first confirmed the antiviral efficacy of NTZ and TIZ *in vitro* before investigating activity against SARS-CoV-2 using reconstituted human airway epithelium and a previously described Syrian hamster model (22, 23). A population pharmacokinetic model was developed to compare exposure in hamsters and humans, with the aim of assessing whether the exposure to NTZ and TIZ in preclinical animal species can be achieved in humans, and whether the antiviral potency observed in vitro can be recovered in vivo.

## Results

### *In vitro* efficacy of nitazoxanide (NTZ)

Using two different cell lines, the VeroE6 (ACE2^+^/TMPRSS2^-^) and Caco-2 cells (ACE2^+^/TMPRSS2^+^), we first evaluated the *in vitro* efficacy of nitazoxanide (NTZ) and tizoxanide (TIZ) against SARS-CoV-2. Anti-viral potency was assessed in a viral RNA yield reduction assay by qRT-PCR as previously described (23-25). In VeroE6 cells, NTZ and TIZ inhibited viral replication with EC_50_’s of 3.19 and 7.48µM and EC_90_’s of 10.27 and 9.27µM, respectively, while both CC_50_’s were above 60µM (Figure 1). Selectivity Index (SI=CC_50_/EC_50_) were higher than 13.3 for both molecules. In Caco-2 cells, NTZ exhibited an EC_50_ of 0.58µM, an EC_90_ of 1.75 µM and a CC_50_ of 9.15µM resulting in a SI of 15.8 (Figure 1).

**Figure 1:**
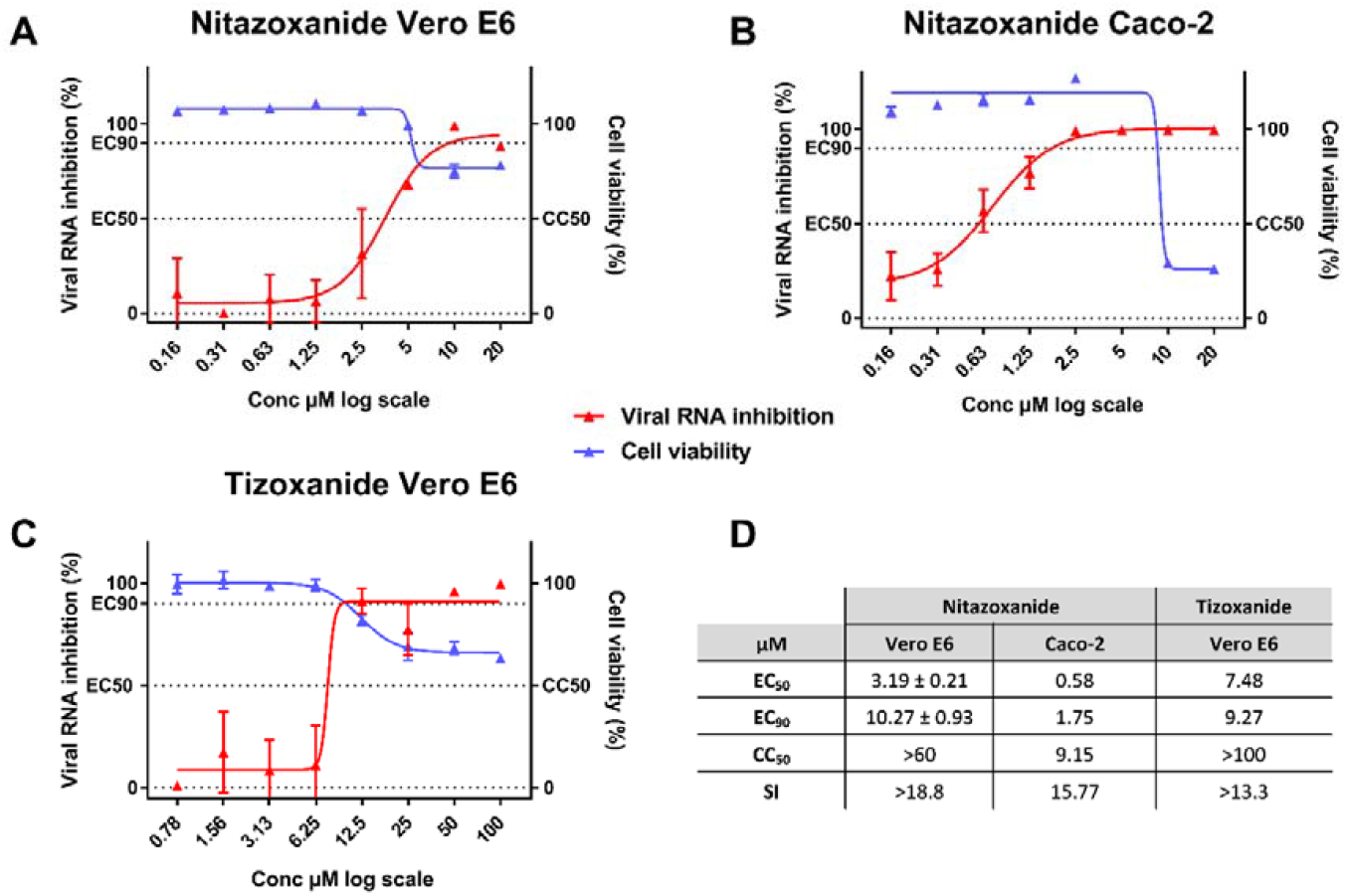
Antiviral activity of NTZ and TIZ in Vero E6 and Caco-2 cells. Dose response curve and cell viability for: NTZ in Vero E6 (a) and Caco-2 (b) cells and for TIZ in Vero E6 cells (c). D: Table of EC_50_, EC_90_, CC_50_. Results presented in the table for NTZ in VeroeE6 are the mean ± SD from three independent experiments. Graphical representation is from one representative experiment.

### *Ex vivo* efficacy of NTZ

We then investigated the *ex vivo* efficacy of NTZ using a recently described model of reconstituted human airway epithelial of bronchial origin (26). Five different concentrations of NTZ (20; 10; 5; 2.5; 1.25µM) were tested in duplicate while Remdesivir, at an active concentration of 10µM, was used as a positive control. The basolateral sides of the epithelia were exposed to the drugs from time of infection until day 4 post infection (dpi). Media with fresh drug were renewed at 1,2 and 3 dpi. Viral excretion was assessed at 2, 3 and 4 dpi, by measuring viral RNA yields and infectious titers at the apical side of the epithelium using quantitative real time RT-PCR and TCID_50_ assays, respectively. No antiviral efficacy was detected when viral excretion was assessed by quantification of viral RNA (Figure 2-A). However, at 3 and 4 dpi, a significant reduction of infectious titers was observed when concentrations of NTZ above its EC_50_ were used (with p values ranging from 0.01-0.05 for 5µM, 0.001-0.01 for 10µM and 0.0001-0.001 for 20µM) (Figure 2).

**Figure 2:**
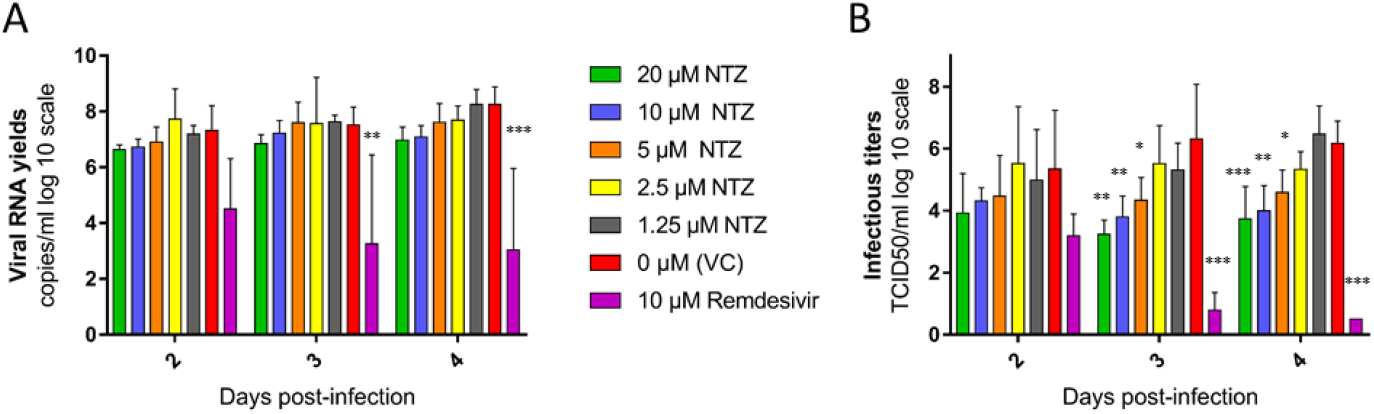
Antiviral activity of NTZ in a bronchial human airway epithelium. Kinetics of virus excretion at the apical side of the epithelium measured using an RT-qPCR assay (A) and a TCID_50_ assay (B). Data represent mean ± SD. Statistical significance was calculated by 1-way ANOVA versus untreated group. Remdesivir at 10µM was used as a positive drug control. *, ** and *** indicate and average significant value lower than that of the untreated group, with a p-value ranging between 0.01-0.05, 0.001-0.01 and 0.0001-0.001, respectively. Result are the mean ± SD of two independent experiment with in each experiment two independent inserts.

### *In vivo* efficacy of NTZ

Considering these results, we further investigated the potential antiviral activity of NTZ *in vivo* using a previously described hamster model of SARS-CoV-2 infection (22, 23, 27).

## Efficacy evaluation of a NTZ oral treatment

In a first set of experiments, we explored the antiviral efficacy of a NTZ suspension (90% sterile distilled water, 7% of tween 80 and 3% of ethanol 80%). During two independent experiments, groups of 6 hamsters were intranasally infected with 10^4^ TCID_50_ of SARS-CoV-2 and received NTZ orally at doses of 500mg/kg/day BID or 750mg/kg/day TID (Figure 3a). Untreated groups of 6 hamsters received the suspension vehicle BID or TID. A group of 6 animals was treated with favipiravir (FVP) intraperitoneally (926mg/kg/day BID) as positive control in one experiment (22).

**Figure 3:**
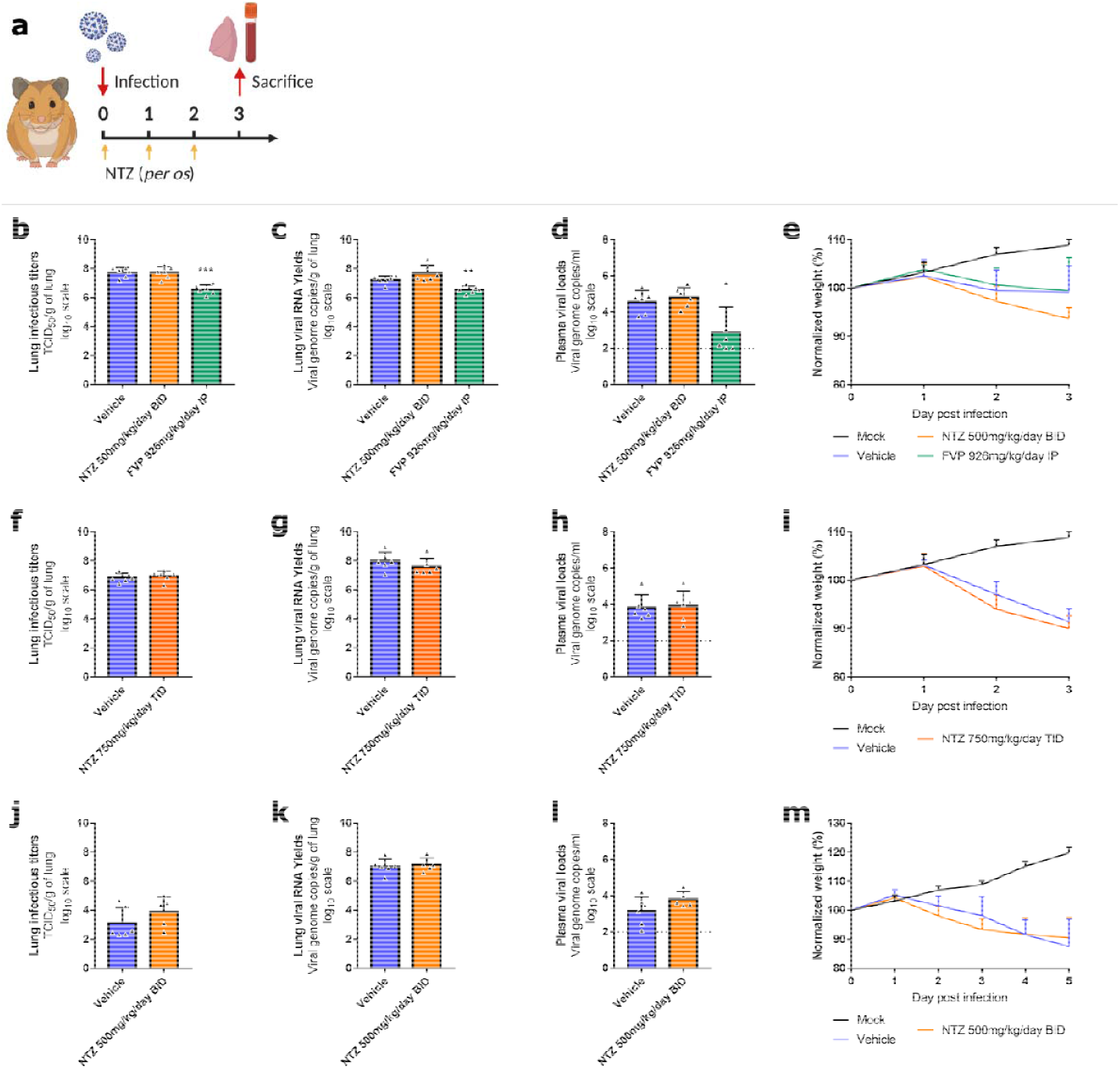
Antiviral activity of oral treatment of NTZ in a hamster model. Groups of 6 hamsters were intranasally infected with 10^4^ TCID_50_ of virus. **a** Experimental timeline. **b, f, j** Viral replication in lung based on infectious titers (measured using a TCID_50_ assay) expressed in TCID_50_/g of lung (n=6 animals/group). **c, g, k** Viral replication in lung based on viral RNA yields (measured using an RT-qPCR assay) expressed in viral genome copies/g of lung (n=6 animals/group). **d, h, l** Plasma viral loads (measured using an RT-qPCR assay) are expressed in viral genome copies/mL of plasma (the dotted line indicates the detection threshold of the assay) (n=6 animals/group). **e, i, m** Clinical course of the disease (n=6 animals/group). Normalized weight at day n was calculated as follows: % of initial weight of the animal at day n. Data represent mean ± SD (Details in Supplementary Data 1). *** and ** symbols indicate that the average value for the group is significantly lower than that of the untreated group with a p-value ranging between 0.0001-0.001 and 0.001-0.01 respectively (Details in Supplementary Data 2).

Hamsters treated with 500mg/kg/day BID or 750mg/kg/day TID of NTZ for 3 days (0, 1 and 2dpi), showed no significant differences for either infectious titers (measured using TCID_50_ assay) or viral RNA yields (measured using quantitative real time RT-PCR assay) in clarified lung homogenates at 3 dpi compared to untreated animals (*p*≥0.0989) (Figure 3b, 3c, 3f and 3g). No significant difference was detected with regards to viral RNA yields in plasma at 3 dpi (*p*≥0.4697) (Figure 3d and 3h). Administration of FVP, however, led as expected to significant reductions of both infectious titers and viral RNA yields in clarified lung homogenates (*p*≤0.0011) (Figure 3b and 3c). NTZ-treated animals showed clinical signs of illness/suffering, with their mean normalized weight becoming significantly lower than that of untreated animals, at 3 dpi for animals treated with 500mg/kg/day BID and at 2 dpi for animals treated with 750mg/kg/day TID (*p*=0.0158 and *p*=0.0314 respectively) (Figure 3e and 3i).

In another independent experiment, we tried to assess the antiviral efficacy of a longer treatment period. Despite receiving 500mg/kg/day of NTZ BID for 4 days (0, 1, 2 and 3 dpi), hamsters exhibited no significant reduction at 5 dpi of either infectious titers (*p*=0.1775) or viral RNA (*p*=0.7003) yields in their clarified lung homogenates, or viral RNA yields plasma (*p*=0.1305) (Figure 3j, 3k and 3l). However, they did not show any clinical signs of illness/suffering compared to untreated animals (Figure 3m).

We also explored the impact of NTZ treatment on lung pathological changes induced by SARS-CoV-2, in an independent experiment. Groups of 4 hamsters, intranasally infected with 10^4^ TCID_50_ of SARS-CoV-2, were orally treated at a dose of 500mg/kg/day BID for 4 days (0, 1, 2 and 3 dpi) (Figure 4a). Untreated hamsters (group of 4 animals) received the suspension vehicle BID. Animals were sacrificed at 5 dpi and a cumulative score from 0 to 10 (taking into account severity of inflammation, alveolar hemorrhagic necrosis and vessel lesions) was calculated and then assigned to a grade of severity (0=normal; 1=mild; 2=moderate; 3=marked and 4=severe; details in Supplementary Data 3 and 4). All animals, treated and untreated, displayed severe pulmonary impairments. Marked and severe histopathological damages in lungs for both groups were identified resulting in no significant difference of histopathological cumulative scores (Figure 4b). At 3 dpi, animals showed clinical signs of illness/suffering, with their mean normalized weight becoming significantly lower than that of untreated animals (Supplementary Fig. 1).

**Figure 4:**
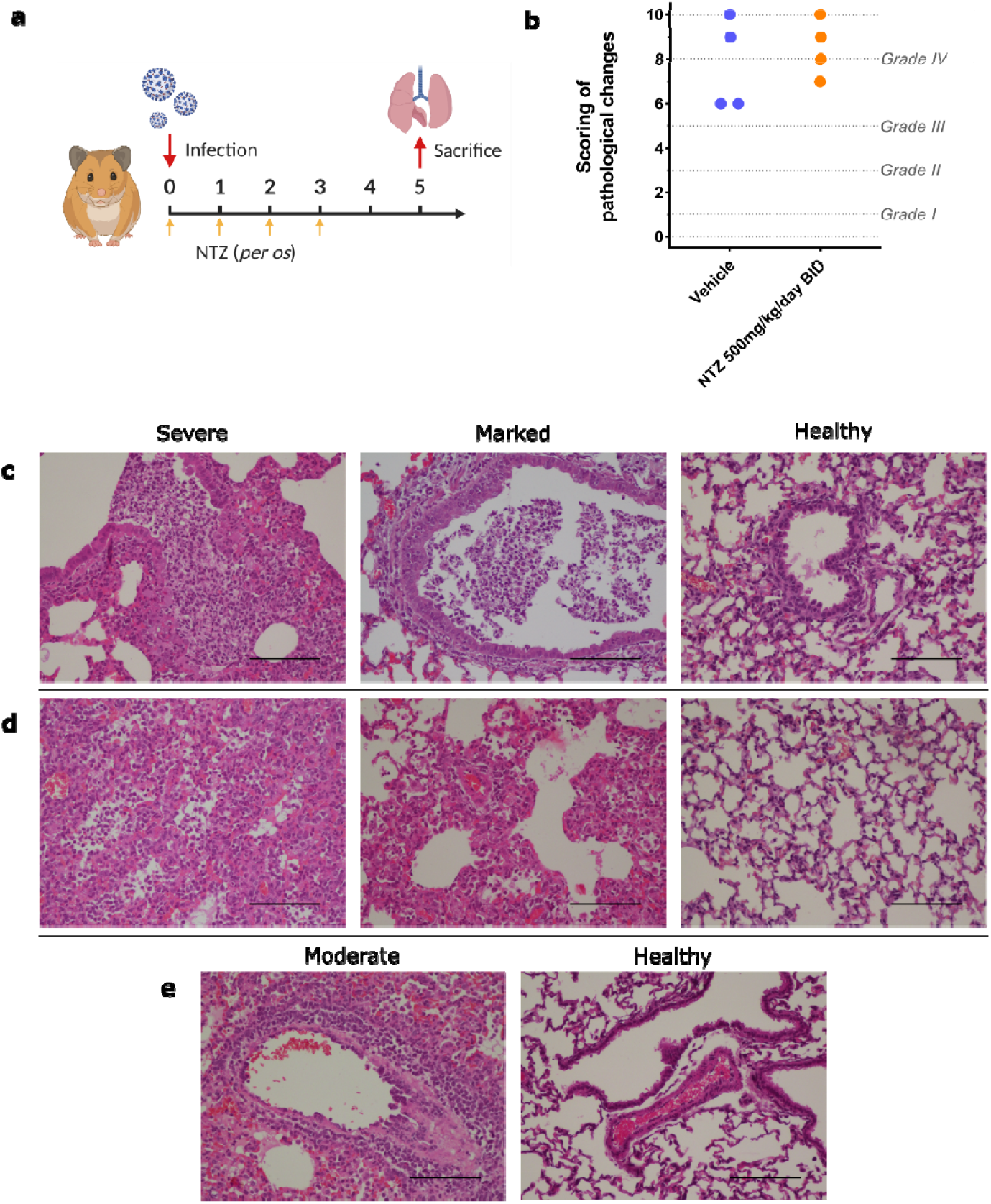
Lung histopathological changes. Groups of 4 animals were intranasally infected with 10 ^4^TCID_50_ of virus and sacrificed at 5 dpi. Based on severity of inflammation, alveolar hemorrhagic necrosis and vessel lesions, a cumulative score from 0 to 10 was calculated and assigned to a grade of severity (I, II, III and IV). **a** Experimental timeline. **b** Scoring of pathological changes (Details in Supplementary Data 3 and 4). **c** Representative images of bronchial inflammation (scale bar: 100µ): severe peribronchiolar inflammation and bronchiole filled with numerous neutrophilic, marked peribronchiolar inflammation and normal bronchi. **d** Representative images of alveolar inflammation (scale bar: 100µ): severe infiltration of alveolar walls, alveoli filled with neutrophils/macrophages, marked infiltration of alveolar walls, some alveoli filled with neutrophils/macrophages and normal alveoli. **e** Representative images of vessel inflammation (scale bar: 100µ): moderate accumulation of inflammatory cells in arteriolar walls and normal arteriole.

To investigate if the lack of efficacy seen in lungs was due to an inadequate drug diffusion, we assessed the exposure and the lung distribution of TIZ (the active circulating metabolite of NTZ). We used tissues from infected animals sacrificed at 3 dpi following multiple administration (animals from Figure 3); an additional group of uninfected animals treated with a single dose of 13.5mg was used as control. TIZ concentration in plasma and in lung was quantified at 1, 2 and 4 hours post treatment for the single dose analysis (group of 3 animals) and at 12 hours after the last administration for the multiple dose analysis (group of 6 animals).

These animals exhibited low penetration rates of TIZ in lungs, resulting in lung/plasma ratio ranging from 2.2% to 4.8% after single-dose administration (Table 1). Lung concentrations of TIZ were below the TIZ EC_50_ found *in vitro* with Vero E6 cells (7.48µM, i.e 1.98µg/mL), as well as effective NTZ concentrations *ex vivo* (5µM, i.e 1.54µg/mL), and were not quantifiable in a total of 5 out of 9 animals (one at 1 hour, two at 2 hours and two at 4 hours) (Table 1). After 3 days of multiple dose treatment, TIZ trough concentrations (12 hours after the last administration) in lungs were still below the *in vitro* EC_50_ and not quantifiable in a total of 8 out of 12 animals (four for each multiple dose concentration) (Table 1).

**Table 1:**
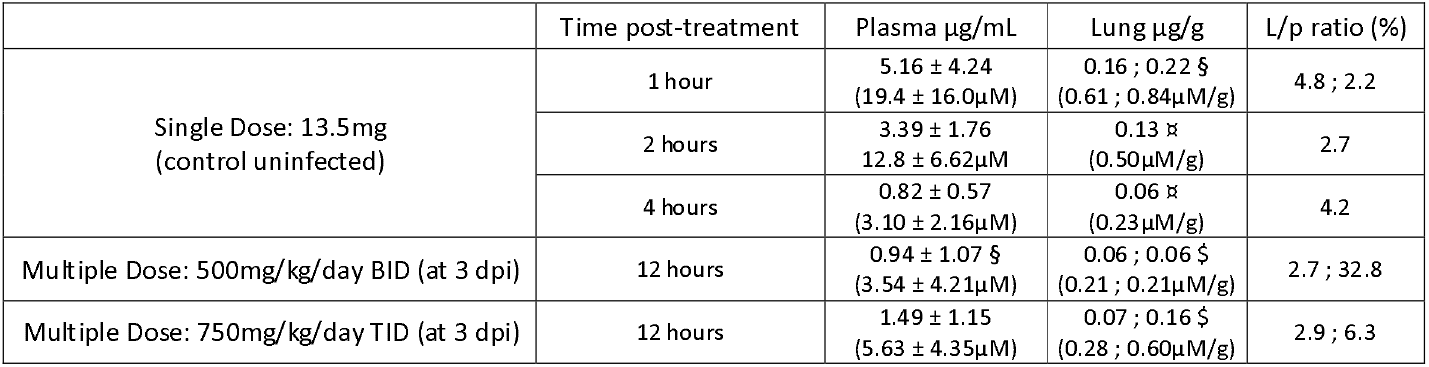
Plasma and lung concentrations of TIZ after administration of a single dose or multiple dose of NTZ. Multiple Dose: PK realized after 3 days of nitazoxanide administered two or three times a day, at the end of the dosing interval (trough concentrations). Data represent mean ± SD for plasma concentrations and individual values for lung concentrations and L/p ratios. These data represent a summary of Supplementary Data 4. Symbols §, ¤ and $ represent respectively 1, 2 and 4 values below the limit of quantification.

### NTZ efficacy evaluation following intranasal administration

To assess other administration routes for NTZ, we explored the antiviral efficacy of an intranasal NTZ emulsion (aqueous phase: sterile distilled water 94% and absolute ethanol 6%; organic phase: NTZ 20mg/mL in cinnamaldehyde 75% and Kolliphore EL 25%). Hamsters were intranasally infected with 10^4^ TCID_50_ of SARS-CoV-2. A group of 6 hamsters received intranasally 2.8mg/kg/day TID of NTZ (Figure 5a). An untreated group of 6 hamsters received the emulsion vehicle TID. All treatments were started at the day of infection and ended at day 2 post infection. Viral replication was assessed in lungs, plasma and nasal turbinates at 3 dpi.

**Figure 5:**
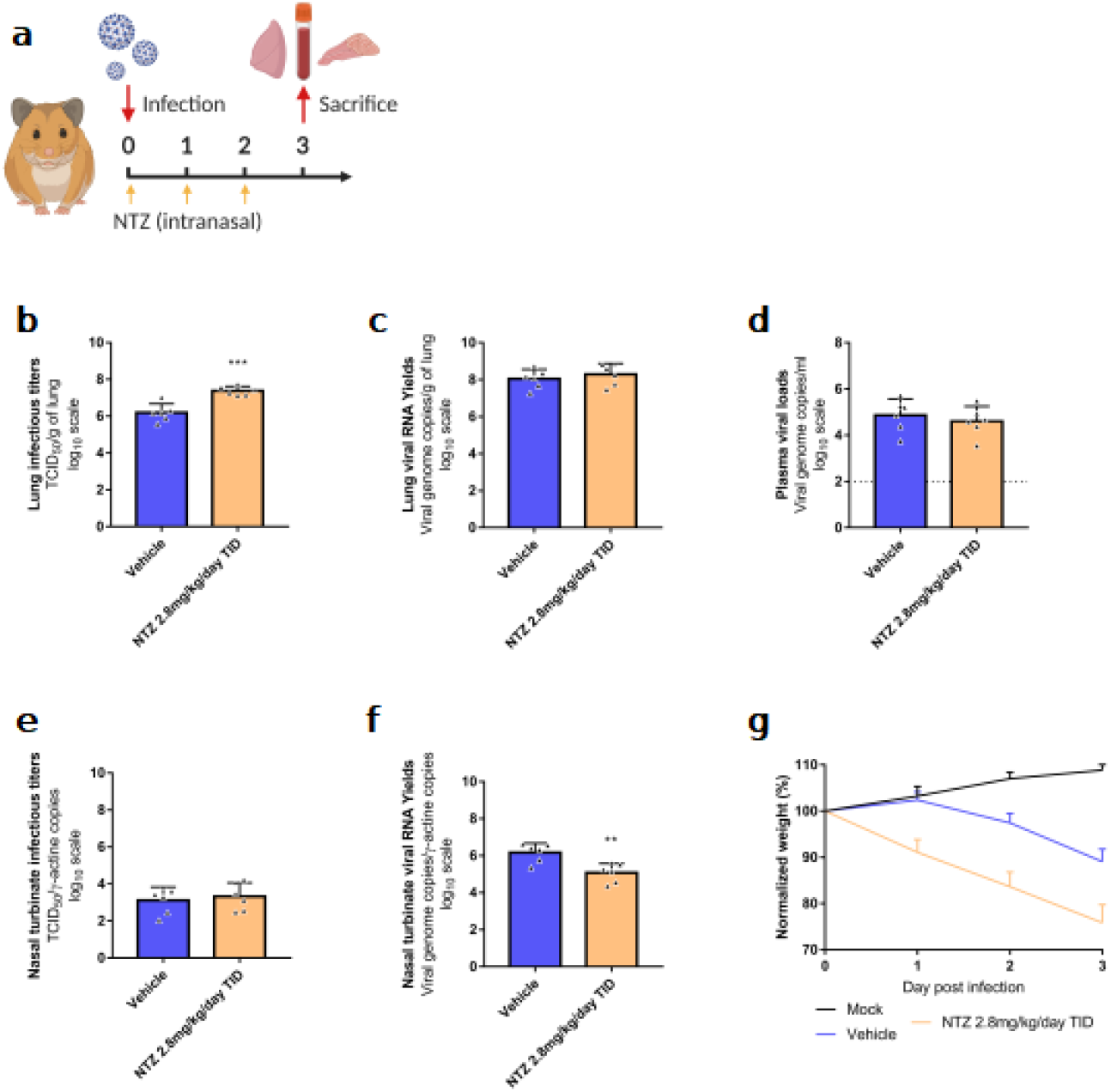
Antiviral activity of intranasal treatment of NTZ in a hamster model. Groups of 6 hamsters were intranasally infected with 10^4^TCID_50_ of virus. **a** Experimental timeline. **b** Viral replication in lung based on infectious titers (measured using a TCID_50_ assay) expressed in TCID_50_/g of lung (n=6 animals/group). **c** Viral replication in lung based on viral RNA yields (measured using an RT-qPCR assay) expressed in viral genome copies/g of lung (n=6 animals/group). **d** Plasma viral loads (measured using an RT-qPCR assay) are expressed in viral genome copies/mL of plasma (the dotted line indicates the detection threshold of the assay) (n=6 animals/group). **e** Viral replication in nasal turbinates based on infectious titers (measured using a TCID_50_ assay) expressed in TCID_50_/copy of ⍰-actine gene (n=6 animals/group). **f** Viral replication in nasal turbinates based on viral RNA yields (measured using an RT-qPCR assay) expressed in viral genome copies/copy of ⍰-actine gene (n=6 animals/group). **g** Clinical course of the disease (n=6 animals/group). Normalized weight at day n was calculated as follows: % of initial weight of the animal at day n. Data represent mean ± SD (Details in Supplementary Data 1). ** symbols indicate that the average value for the group is significantly lower than that of the untreated group with a p-value ranging between 0.001-0.01 (Details in Supplementary Data 1).

**Figure 6:**
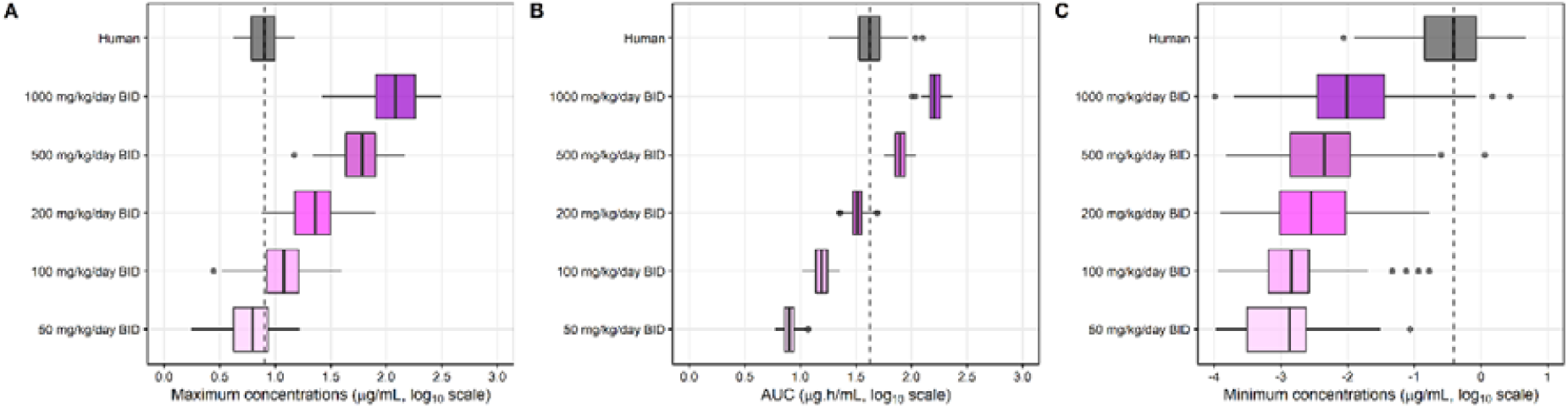
Simulated pharmacokinetic parameters of TIZ in human and hamster at steady state. Predicted steady-state pharmacokinetic parameters of TIZ, i.e. C_max_ (A), AUC (B) and C_min_ (C), in human associated with receiving 1000mg/day BID of NTZ (grey box) were compared with pharmacokinetic parameters of hamster receiving 50, 100, 200, 500 and 1000mg/kg/day BID of NTZ (colored boxes). Boxes and whiskers represent the median with inter-quantile range and the 95% prediction intervals, respectively.

NTZ intranasal treatment led to a significant increase of infectious titers in clarified lung homogenates (*p*=0.0003) (Figure 5b). No significant differences were observed when looking at viral RNA yields in both clarified lung homogenates and plasma with both intranasal treatments (*p*≥0.4530) (Figure 5c and 5d). In nasal turbinates, no significant differences of infectious titers were observed between the groups (*p*=0.6295) (Figure 5e). When looking at viral RNA yields, NTZ treatment induced a significant reduction of viral RNA load in nasal turbinates (*p*=0.0037) (Figure 5f). Animals treated with NTZ intranasally from 1 to 3 dpi, showed clinical signs of illness/suffering, with their mean normalized weight becoming significantly lower than that of untreated animals (*p*=0.0314) (Figure 5g).

To confirm these results, the experiment was repeated independently. Overall, no significant differences in viral replication between treated and untreated hamsters were observed in either clarified lung homogenates, plasma or nasal turbinates (Supplementary Fig. 2). Once again, hamsters treated with NTZ intranasally from 1 to 3 dpi, showed clinical signs of illness/suffering, with their mean normalized weight becoming significantly lower than that of untreated animals (Supplementary Fig. 2).

Similarly to the oral administration study, and to investigate a potential issue regarding drug distribution to the compartment of choice, we also assessed the plasma, lung and nasal turbinates concentrations of TIZ following intranasal NTZ administration in infected animals treated by multiple doses, 12 hours after the last administration (group of 6 hamsters from figure 5). Overall, TIZ was detectable in only 2 out of 6 animals (one in plasma and lung; one in lung and nasal turbinates) (Supplementary Table 1) but TIZ concentrations were below the EC_50_ found *in vitro* and the active concentration *ex vivo*.

### Pharmacokinetic modelling

We characterized the pharmacokinetic profile of TIZ in hamster after administration of the same NTZ suspension or TIZ formulated in 10% [Tween 80, 80% EtOH (70:30 v/v)] and 90% distilled water, homogenous opaque suspension. Groups of 3 hamsters received an oral single dose of 485mg/kg, 98.1mg/kg or 25.5mg/kg of NTZ or 96.4mg/kg of TIZ. The corresponding concentration-time curves for NTZ administration only are presented in Supplementary Fig. 3. Notably, similar concentration-time curves of TIZ were observed following oral administration of 98.1 NTZ or 96.4mg/kg of TIZ, suggesting full in-vivo conversion of NTZ into TIZ (Supplementary Fig. 3). The observed TIZ plasma concentration-time data in hamster, following oral administration of NTZ and TIZ, were characterized using nonlinear mixed-effects modelling. The data were best described by a two-compartment disposition model with first-order absorption. The population pharmacokinetic parameters estimates from the final model are presented in Supplementary Table 2. Time to maximum concentration (T_max_) and terminal elimination half-life (t_1/2_) were estimated to 0.276h and 0.80h, respectively. Graphically, the model showed good adequacy between predicted concentrations and observed concentrations (Supplementary Fig. 4)

Simulated secondary pharmacokinetic parameters (C_max_ and AUC_0-24h_) at each NTZ dose derived from the population pharmacokinetic model are presented in Table 2.

**Table 2:** Simulated pharmacokinetic parameters derived from nonlinear mixed-effect modelling using population mean values and median body weight of 0.126kg. C_max_: maximum plasma drug concentration; AUC_0-24h_: area under the concentration-time curve from time 0 to 24h.

| NTZ dose (mg/kg) | $C_{max}$ (μg/ml) | $AUC_{0-24h}$ (μg·h/ml) |
| --- | --- | --- |
| 25.0 | 3.14 | 4.18 |
| 50.0 | 6.27 | 8.36 |
| 100 | 12.5 | 16.7 |
| 125 | 15.7 | 20.9 |
| 250 | 31.4 | 41.8 |
| 500 | 62.7 | 83.6 |

We then compared the pharmacokinetic profile of TIZ in hamster to that in human. The pharmacokinetic model developed to describe the TIZ concentration-time data in hamster was used to simulate plasma drug exposures in hamster using different dose regimens. From these simulations, TIZ C_max_, AUC (AUC at steady state over 1 dosing interval) and C_min_ were derived and compared to simulated plasma PK parameters in human using a published physiologically-based pharmacokinetic (PBPK) model (28). A dose between 50 and 100mg/kg/day BID in hamsters provided C_max_ values close to that obtained in humans after a dose of 1000mg BID. A dose between 200 and 500mg/kg BID in hamster provided AUC values quite similar to the one obtained in humans. However, human C_min_ was never reached at any dose in hamsters.

## Discussion

Nitazoxanide was among the very first molecules studied at the beginning of the COVID-19 pandemic, revealing an *in vitro* antiviral efficacy against SARS-CoV-2 (13-19). Our study confirms these results, as we found that NTZ possesses EC_50_ under 5µM in two different cell lines. In addition, we demonstrated that NTZ was active in bronchial human airway epithelia, which largely mimic the structural, functional, and innate immune features of the human respiratory epithelium, albeit at lower potency as compared to Remdesivir, the positive control in this assay (26).

It is well documented that, *in vivo*, NTZ is rapidly deacetylated to its active metabolite, TIZ (29). However, only one non-peer-reviewed source reported the activity of TIZ against SARS-CoV-2 (https://opendata.ncats.nih.gov/covid19/databrowser). Here, we demonstrated that this metabolite is indeed active against SARS-CoV-2 *in vitro* with an EC_50_ of 7.48µM, reinforcing the potential use of NTZ in COVID-19 management.

Although mainly considered as an antiprotozoal agent, NTZ, and its active circulating TIZ metabolite, were identified as *in vitro* broad-spectrum antiviral compounds, since they both inhibit the replication of a wide range of RNA and DNA viruses in cell culture (6, 8). Their antiviral mechanism has not been clearly elucidated. However, it seems that interaction with numerous targets implicated in viral pathogenesis, depending on the virus, is involved (immunomodulatory effects and direct drug action) (8). Notably, post-entry inhibition by upregulation of cell’s innate antiviral response, observed against hepatitis C virus on cell culture (30), may be one of the efficient antiviral pathways involved in SARS-CoV-2 inhibition. Recently, TMEM16 inhibitors, such as niclosamide and NTZ, have been reported to protect against cell fusion induced by SARS-CoV-2 spike protein in cell culture (31). This syncytia inhibition could be one of the modes of action observed *in vitro* and *ex vivo* for NTZ antiviral activity against SARS-CoV-2.

Although NTZ has had only incomplete preclinical characterization, numerous clinical trials using the molecule are underway around the world. We felt that the generation of robust preclinical data was relevant to document the suitability of NTZ for clinical use in the treatment of COVID patients.

Despite promising *in vitro* results and new hypotheses on its antiviral mechanism, NTZ failed to reduce the severity of SARS-CoV-2 infection *in vivo* in the Syrian hamster model. No significant improvement in terms of clinical course of the disease, viral replication (based on infectious titers or viral RNA yields) and/or histopathological damages in lungs was observed when using two different dosing regimens of NTZ. These findings could be explained by the insufficient pulmonary diffusion of TIZ, since peak concentrations in lungs (1 hour post-treatment) never exceeded its *in vitro* or *ex vivo* EC_50_. This insufficient pulmonary exposure was confirmed by the low accumulation of TIZ over time in the lungs as trough concentrations after 3 days of multiple doses of NTZ were similar to those found 4 hours post-treatment in the single dose model. This result is in accordance with a previous clinical trial assessing the safety, bactericidal activity, and pharmacokinetics of NTZ in adults with pulmonary tuberculosis, where sputum concentration of NTZ was low, suggesting that it did not penetrate pulmonary lesions to a sufficient degree (32). This can be explained in part by the physico-chemical and pharmacokinetic characteristics of the product, which do not facilitate tissue diffusion. In addition to being a moderately lipophilic molecule, TIZ is highly bound to plasma proteins (99%). Therefore, the use of NTZ as a systemic treatment might be challenging.

The PK modeling and simulations provided further insights for the lack of NTZ efficacy in the *in vivo* hamster model of SARS-CoV-2 infection. Simulations showed that the dose of 500mg/kg/day BID (found ineffective in our study) was sufficient to achieve C_max_ and AUC above those observed in humans at the usual dose of 1000mg/kg/day, but not sufficient to reach trough concentrations (C_min_) observed in humans at this same dose. Furthermore, human C_min_ was never reached even with the highest simulated dose (1000mg/kg/day BID). This prediction was confirmed in our *in vivo* study in which we did not observe efficacy at the highest dose (*i*.*e*. 750mg/kg/day TID). The latter observations show a difference in the clearance of NTZ between humans and hamsters, of which the C_min_ can be considered a reflection, with a more rapid elimination in hamsters. Although differences in pharmacokinetic profiles between humans and hamsters are known and widely documented, these findings suggest that at the usual dose of 1000mg/kg/day in humans, NTZ will have no effect on SARS-CoV-2 replication.

To potentially enhance the pulmonary diffusion and explore the possible antiviral activity of TIZ within the upper respiratory tract, hamsters were treated with an intranasal NTZ emulsion formulation. This alternative route of administration proved ineffective in our model, as no significant improvement in any of the disease endpoints analysed was observed. As observed after oral administration, TIZ trough concentration measured in lungs after 3 days of intranasal NTZ administration was very low, which may also partly explain the lack of antiviral efficacy; in all samples tested, including nasal turbinates, TIZ concentration was found to be well below the *in vitro* and *ex vivo* EC_50_. This lack of TIZ accumulation in the upper respiratory tract should be interpreted with caution as no active intranasal deliverable compounds was available as a positive control in our study.

Overall, based on the pharmacokinetic data collected in this pre-clinical study, the use of NTZ as an antiviral against SARS-CoV-2, does not seem appropriate at the current standard formulation and dosage. Our results suggest that the low pulmonary bioavailability of NTZ remains the major challenge that needs to be addressed in order to properly evaluate the potential antiviral effect of NTZ in an animal model or in human.

Clinical trials with NTZ are currently ongoing and their outcome will be very useful for back-translation purposes. As an example, if preliminary data of a recent trial may suggest that NTZ could have some beneficial impact in preventing worsening of the disease and need for hospitalization, qualitative and quantitative tests to detect SARS-CoV-2 were not significantly different between the treatment arms (33). These observations corroborate our results and demonstrate that it will be essential to increase the pulmonary bioavailability of NTZ in order to conclude a direct antiviral impact.

In conclusion, optimization of the NTZ formulation may allow reconsideration of the potential use of the drug for the treatment of SARS-CoV-2 infection. In a previous pharmacokinetic study of NTZ in mice, optimal concentrations of TIZ were obtained in the lungs when the molecule was entrapped in inhalable particles (34). This type of formulation combined with aerosol administration could potentially lead to an effective concentration of NTZ in the animal’s lungs and deserves further investigation.

## Methods

### Cells and human airway epithelia

VeroE6 cells (ATCC CRL-1586) and Caco-2 cells (ATCC HTB-37) were cultivated under 5% CO_2_ and at 37.5°C in minimal essential medium (MEM) supplemented with 7.5% heat-inactivated fetal bovine serum (FBS), 1% non-essential amino acids and 1% Penicillin/Streptomycin (all from Life Technologies).

Mucilair™ human airway epithelia (HAE), reconstituted from primary cells of bronchial biopsies of a 56-year-old donor Caucasian female with no reported pathologies, was maintained in air liquid interface with specific media (all from Epithelix SARL, Geneva, Switzerland, with informed consent).

### Virus

SARS-CoV-2 strain BavPat1 was provided by Pr.Christian Drosten (Berlin, Germany) through European Virus Archive GLOBAL (https://www.european-virus-archive.com/). Inoculation with this strain at a MOI of 0.001, of a 25cm^2^ culture flask of confluent VeroE6 cells with MEM medium supplemented with 2.5% FBS, allowed us to prepare virus working stocks. Each 24h the cell supernatant medium was replaced in order to be harvested at the peak of infection. It was supplemented with 25mM HEPES (Sigma-Aldrich), aliquoted and stored at -80°C. Experiments with infectious virus were performed in a biosafety level 3 laboratory.

### *In vitro* determination of EC_50_ and CC_50_

One day prior to infection, 96-well culture plates were seeded with 5×10^4^ VeroE6 or Caco-2 cells in 100µL assay medium per well (containing 2.5% FCS). The next day, eight 2-fold serial dilutions of compounds (from 20µM to 0.16µM for NTZ (BLDpharm) and from 100µM to 0.78µM for TIZ (MedChemExpress)) in triplicate were added to the cells (25µL/well, in assay medium). For the determination of the 50% and 90% effective concentrations (EC_50_, EC_90_; compound concentration required to inhibit by 50% or 90% viral RNA replication), four “virus control” wells were supplemented with 25µL of assay medium without any compounds. After 15min, a preset amount of virus diluted in 25µL of assay medium was added to the wells. This quantity of virus was calibrated so that the viral replication was still in the exponential growth phase for the readout, as previously described (24, 25, 35). Four “cell control” wells were supplemented with 50µL of assay medium without any compounds or virus. On each culture plate, a positive control compound (Remdesivir, BLDpharm) was added in duplicate with eight 2-fold serial dilutions (0.16µM to 20µM). Plates were incubated for 2 days at 37°C prior to quantification of the viral genome by real-time RT-PCR as described below. For the determination of the 50% cytotoxic concentrations (CC_50_; compound concentration required to reduce by 50% cell viability), the same culture conditions were used, without addition of the virus, and cell viability was measured using CellTiter Blue® (Promega) following manufacturer’s instructions. EC_50_, EC_90_ and CC_50_ were determined using logarithmic interpolation as previously described (25). The selectivity index of the compounds was calculated as the ratio of the CC_50_ over the EC_50_.

### *Ex vivo* determination of antiviral activity

After being washed with pre-warmed OptiMEM medium (Life technologies), human airway epithelia were infected with SARS-CoV-2 at the apical side using a MOI of 0.1, as previously described (Pizzorno et al., 2020). Cells were cultivated in a basolateral medium that contained NTZ or remdesivir (positive control) at different concentrations or with no drug (virus control). Each day, medium was renewed and samples containing viral RNA were collected by washing the apical side with 200µL of pre-warmed OptiMEM medium. Four day after the infection, total intracellular RNA of each well was extracted using the RNeasy 96 HT kit (Qiagen) following manufacturer’s instructions. Viral RNA was quantified by RT-qPCR and infectious titers were determined in daily samples by TCID_50_, both described below. *Ex vivo* experiments were approved by ethical committee and were conducted according to the declaration of Helsinki on biomedical research (Hong Kong amendment, 1989).

### *In vivo* experiments

#### Approval and authorization

*In vivo* experiments were approved by the local ethical committee (C2EA—14) and the French ‘Ministère de l’Enseignement Supérieur, de la Recherche et de l’Innovation’ (APAFIS#23975). Animal experimentations were performed in accordance with the French national guidelines and the European legislation covering the use of animals for scientific purposes.

#### Animal handling

Three-week-old female Syrian hamsters were provided by Janvier Labs. Animals were maintained in ISOcage P -Bioexclusion System (Techniplast) with unlimited access to water/food and 14h/10h light/dark cycle. Animals were weighed and monitored daily for the duration of the study to detect the appearance of any clinical signs of illness/suffering. General anesthesia was obtained with isoflurane (Isoflurin®, Axience). Euthanasia, which was also realized under general anesthesia, was performed by cervical dislocation.

#### Hamster Infection

Four-week-old anesthetized animals were intranasally infected with 50µL containing 10^4^ TCID_50_ of virus in 0.9% sodium chloride solution. The mock-infected group was intranasally inoculated with 50µL of 0.9% sodium chloride solution.

#### Drug preparation and administration

Hamsters were orally treated with either a NTZ solution at 10mg/mL, suspension at 27mg/mL or emulsion at 2.5mg/mL, prepared from NTZ powder (BLD Pharm). The solution was prepared with 0.5% of hydroxypropyl methylcellulose and 0.1% of tween 80. For the suspension NTZ was dissolved in a vehicle composed of 90% (v / v) sterile distilled water, 7% (v / v) of tween 80 and 3% (v / v) ethanol 80%. The emulsion (aqueous/organic phase ratio of 80/20) for intranasal instillation was prepared with an aqueous phase (sterile distilled water 94% and absolute ethanol 6%) added gradually to an organic phase (NTZ 20mg/mL in cinnamaldehyde 75% and Kolliphore EL 25%) under constant stirring. A solution of favipiravir, reconstituted from anhydrous favipiravir (Toyama-Chemical) with 0.9% sodium chloride solution, was used for intra-peritoneally and intranasally treatment. Control group were orally or intranasally inoculated with a 0.9% sodium chloride solution.

#### Tissue collection

Lungs, nasal turbinates and blood were collected immediately after euthanasia. The left pulmonary lobe was first rinsed in 10mL of 0.9% sodium chloride solution, blotted with filter paper and weighed. Nasal turbinates and pulmonary lobes were transferred to a 2mL tube containing respectively 500µL or 1mL of 0.9% sodium chloride solution and 1mm or 3mm glass beads. They were crushed using a Tissue Lyser machine (Retsch MM400) for 5min at 30 cycles/s and then centrifuged 10min at 16,200g. Crushed nasal tubinates were stored at -80°C while lung supernatant media were transferred to a 1.5mL tube, for another centrifugation during 10 min at 16,200g prior being stored at -80°C. One milliliter of blood was harvested in a 2mL tube containing 100µL of 0.5M EDTA (Life Technologies). Blood was centrifuged for 10 min at 16,200g and stored at -80°C.

### Quantitative real-time RT-PCR (RT-qPCR) assays

All experiments were conducted in a molecular biology laboratory that is specifically devoted to molecular clinical diagnosis and which includes separate laboratories dedicated to each step of the procedure. Prior to PCR amplification, RNA extraction was carried out using the QIAamp 96 DNA kit and the Qiacube HT kit and the Qiacube HT (both from Qiagen) following the manufacturer’s instructions. Shortly, 100µl of tissue clarified homogenates, spiked with 10µL of internal control (bacteriophage MS2), or viral supernatant were transferred into an S-block containing the recommended volumes of VXL, proteinase K and RNA carrier.

RT-qPCR (SARS-CoV-2 and MS2 viral genome detection) were performed with the GoTaq 1-step qRt-PCR kit (Promega) using 3.8µL of extracted RNA and 6.2µL of RT-qPCR mix that contains 250nM of each primer and 75nM of probe. Primers and probes sequences used are described in Supplementary Table 3. Quantification was provided by four 2 log serial dilutions of an appropriate T7-generated synthetic RNA standard of known quantities (10^2^ to 10^8^ copies/reaction). Amplification was performed with the QuantStudio 12K Flex Real-Time PCR System (Applied Biosystems) using standard fast cycling parameters: 10min at 50°C, 2 min at 95°C, and 40 amplification cycles (95°C for 3 sec followed by 30sec at 60°C). qPCR (1-actine gene detection) was performed under the same condition as RT-qPCR with the following modifications: we used the Express one step qPCR Universal kit (ThermoFisher Scientific) and the 50°C step of the amplification cycle was removed. Results were analyzed using QuantStudio 12K Flex Applied Biosystems software v1.2.3.

### Tissue-culture infectious dose 50 (TCID_50_) assay

To determine infectious titers, 96-well culture plates containing confluent VeroE6 cells were inoculated with 150μL per well of serial dilutions of each sample (ten-fold or four-fold dilutions when analyzing cell supernatant media or lung clarified homogenates respectively). Each dilution was performed in sextuplicate. After 5 days of incubation, plates were read for the absence or presence of cytopathic effect in each well. Infectious titers were estimated using the method characterized by Reed & Muench (36).

### Nitazoxanide quantification in plasma and tissues

Quantification of TIZ in plasma and lung tissues was performed by high-performance liquid chromatography with UV detection method (Alliance 2695, Waters, USA) with a lower limit of quantification of 0.01µg/mL. The mobile phase consisted of 0.1% FA in water and 0.1% of FA in ACN (65:35, v/v). The chromatographic separation was achieved using an isocratic mode with an Xbridge BEH C18 2.5 μm 4.6 × 100mm column. Peak area was quantified at 340nm using the Waters 2489 detector. TIZ was extracted by a simple protein precipitation method, using acetonitrile for plasma and ice-cold acetonitrile for clarified lung homogenates. Briefly, 200µL of samples matrix was added to 1000µL of acetonitrile solution containing the internal standard (thiopental), then vortexed for 2min followed by centrifugation for 10min at 4°C. The supernatant medium was evaporated under vacuum, then transferred to a 1.5mL Eppendorf tube. The dried residue was reconstituted with 100µl of ACN:water (50:50), vortexed for 30 seconds and centrifuged again for 10min at 4°C. The supernatant was transferred to an autosampler and 50µL was injected.

### Histology

Animal handling, hamster infections, NTZ preparation and oral administrations were performed as described above. The anatomo-histological study was implemented as previously described (22). Briefly, lungs were collected after intratracheal instillation of 4% (w/v) formaldehyde solution, and then fixed 72h at room temperature with a 4% (w/v) formaldehyde solution before being embedded in paraffin. Tissue sections of 3.5µm were stained with hematoxylin-eosin (H&E) and blindly analyzed by a certified veterinary pathologist. Microscopic examination was done using a Nikon Eclipse E400 microscope. Different anatomic compartments were examined (1) for bronchial and alveolar walls, a score of 0 to 4 was assigned based on severity of inflammation; (2) regarding alveoli, a score of 0 to 2 was assigned based on presence and severity of hemorrhagic necrosis; (3) regarding vessel changes (leucocytic accumulation in vascular wall or in endothelial compartment), absence or presence was scored 0 or 1 respectively. A cumulative score was then calculated and assigned to a grade of severity (see Supplementary Table 4).

### Pharmacokinetic modelling and simulation

NTZ is rapidly and completely hydrolyzed into its active metabolite TIZ (29, 37). Therefore, the pharmacokinetic properties of NTZ were described using measured TIZ concentration in plasma. At each time point, approximate 80µl of blood were collected from the submandibular vein or the saphenous vein of hamsters. All samples were transferred into commercial K2-EDTA tubes, placed on ice until processed for plasma extraction by centrifugation and stored at -70°C before analysis. A LC-MS/MS-AI Triple Quad 5500 was used to determine TIZ concentrations. The mobile phase was a gradient of 0.1% formic acid (FA) in water and 0.1% of FA in acetonitrile (ACN), the column was an ACQUITY UPLC HSS T3 1.8μm 2.1 × 50mm. For mass spectrometry a positive electrospray ionization was used, and a selected reaction monitoring was set to select TIZ: [M+H]+m/z: 266.0 / 121.2 and dexamethasone: [M+H]+m/z: 393.0 / 373.1 as internal standard.

Pharmacokinetic profiles of TIZ in hamster were analyzed using a nonlinear mixed-effects modelling approach. The population pharmacokinetics analysis was performed using NONMEM® version 7.4. The final pharmacokinetic model was established by evaluating one-, two-, and three-compartment disposition models, as well as several different absorption models (i.e. first-order absorption, first-order absorption with lag time, and transit absorption models). The inter-individual variabilities of pharmacokinetic parameters were implemented as a log-normal distribution and the residual unexplained variability was modelled as an exponential error. Different doses of NTZ (i.e., 50, 100, 200, 500 and 1000mg/kg/day BID) were used to simulate TIZ exposures in hamsters (n=100) in order to compare to the simulated human exposure.

To simulate human PK profiles, a one-compartment model was used with pharmacokinetic parameters from an established PBPK model (28), developed to describe pharmacokinetic data of TIZ plasma concentrations in healthy individuals receiving single doses of 500-4000mg NTZ with/without food, presenting an apparent clearance of 19.34L/h and a volume of distribution of 38.68 L. The absorption was described with a first-order process and the rate constant (k_a_) of 0.45h^-1^ was assumed in order to generate the mean concentration-time profile with a T_max_ at approximately 2 hours, as reported in healthy volunteers (38). This model was used to simulate drug exposure at steady state in human after a dosing regimen of 1000mg/day BID of NTZ, administered to 100 individuals. A total of 10 doses (5 days) were used for estimating the convergence rate to steady state.

All exposure simulations were performed in R version 4.0.4 using the mlxR package (39).

### Graphical representations and statistical analysis

Graphical representations and statistical analyses (two-sided tests when relevant) were performed with Graphpad Prism 7 (Graphpad software). P-values lower than 0.05 were considered statistically significant. Statistical details for each experiment are described in the figure legends and in corresponding Supplementary Data. Experimental timelines were created on biorender.com.

## Acknowledgments

We thank Laurence Thirion (UVE; Marseille) for providing RT-qPCR systems. We thank Camille Placidi (UVE; Marseille) for her technical contribution. We thank Pr Drosten and Pr Drexler for providing the SARS-CoV-2 strain through the European Research infrastructure EVA GLOBAL. We thank Toyama-Chemical Favipiravir for kindly providing the favipiravir. This work was supported by the Fondation de France “call FLASH COVID-19”, project TAMAC, by “Institut national de la santé et de la recherche médicale” through the REACTing (REsearch and ACTion targeting emerging infectious diseases), by REACTING/ANRS MIE under the agreement No. 21180 (‘Activité des molécules antivirales dans le modèle hamster’), by European Virus Archive Global (EVA 213 GLOBAL) funded by the European Union’s Horizon 2020 research and innovation program under grant agreement No. 871029 and DND*i* under support by the Wellcome Trust Grant ref: 222489/Z/21/Z through the COVID-19 Therapeutics Accelerator”. Part of this work was supported by the Wellcome Trust [220211]. Part of the work was done on the Aix Marseille University antivirals platform “AD2P”. For the purpose of open access, the author has applied a CC BY public copyright licence to any Author Accepted Manuscript version arising from this submission.

## Supplementary Data

**Supplementary Fig. 1:**
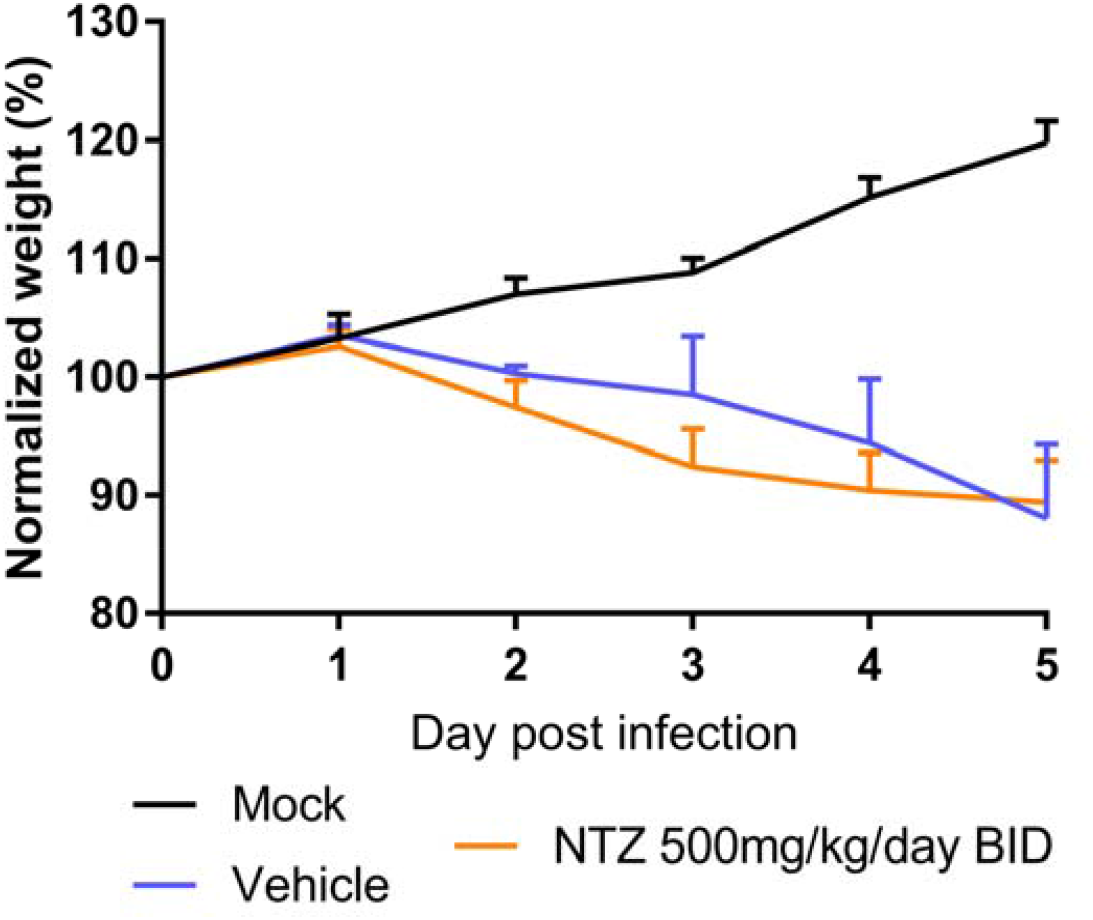
Clinical course of the disease (n=4 animals/group). Normalized weight at day n was calculated as follows: % of initial weight of the animal at day n. Data represent mean ± SD (Details in Supplementary Data 3).

**Supplementary Fig. 2:**
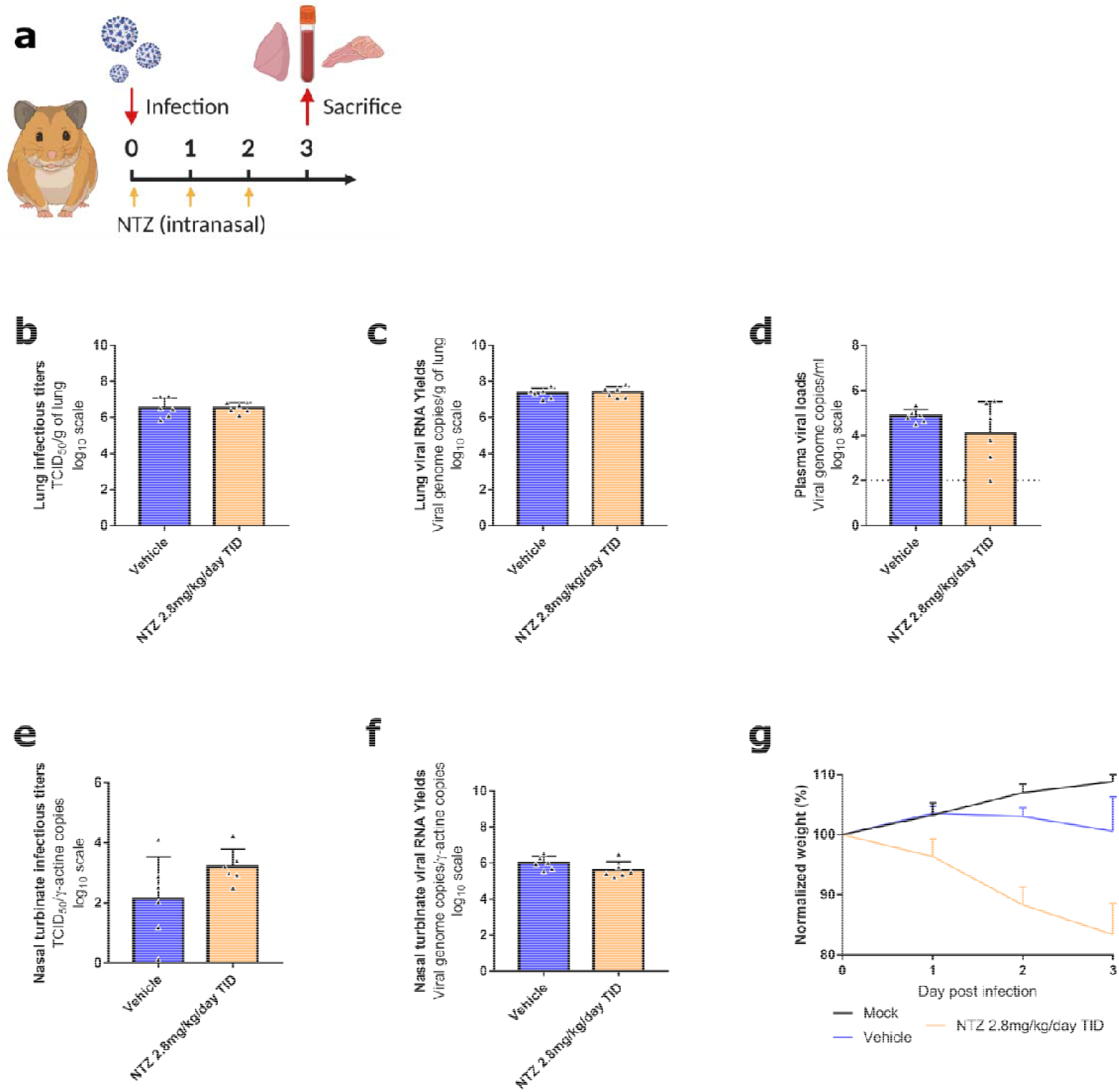
Antiviral activity of intranasal treatment of NTZ in a hamster model. Groups of 6 hamsters were intranasally infected with 10^4^ TCID_50_ of virus. **a** Experimental timeline. **b** Viral replication in lung based on infectious titers (measured using a TCID_50_ assay) expressed in TCID_50_/g of lung (n=6 animals/group). **c** Viral replication in lung based on viral RNA yields (measured using an RT-qPCR assay) expressed in viral genome copies/g of lung (n=6 animals/group). **d** Plasma viral loads (measured using an RT-qPCR assay) are expressed in viral genome copies/mL of plasma (the dotted line indicates the detection threshold of the assay) (n=6 animals/group). **e** Viral replication in nasal turbinates based on infectious titers (measured using a TCID_50_ assay) expressed in TCID_50_/copy of ⍰-actine gene (n=6 animals/group). **f** Viral replication in nasal turbinates based on viral RNA yields (measured using an RT-qPCR assay) expressed in viral genome copies/copy of ⍰-actine gene (n=6 animals/group). **g** Clinical course of the disease (n=6 animals/group). Normalized weight at day n was calculated as follows: % of initial weight of the animal at day n. Data represent mean ± SD (Details in Supplementary Data 1). ** symbols indicate that the average value for the group is significantly lower than that of the untreated group with a p-value ranging between 0.001-0.01 (Details in Supplementary Data 1).

**Supplementary Fig. 3:**
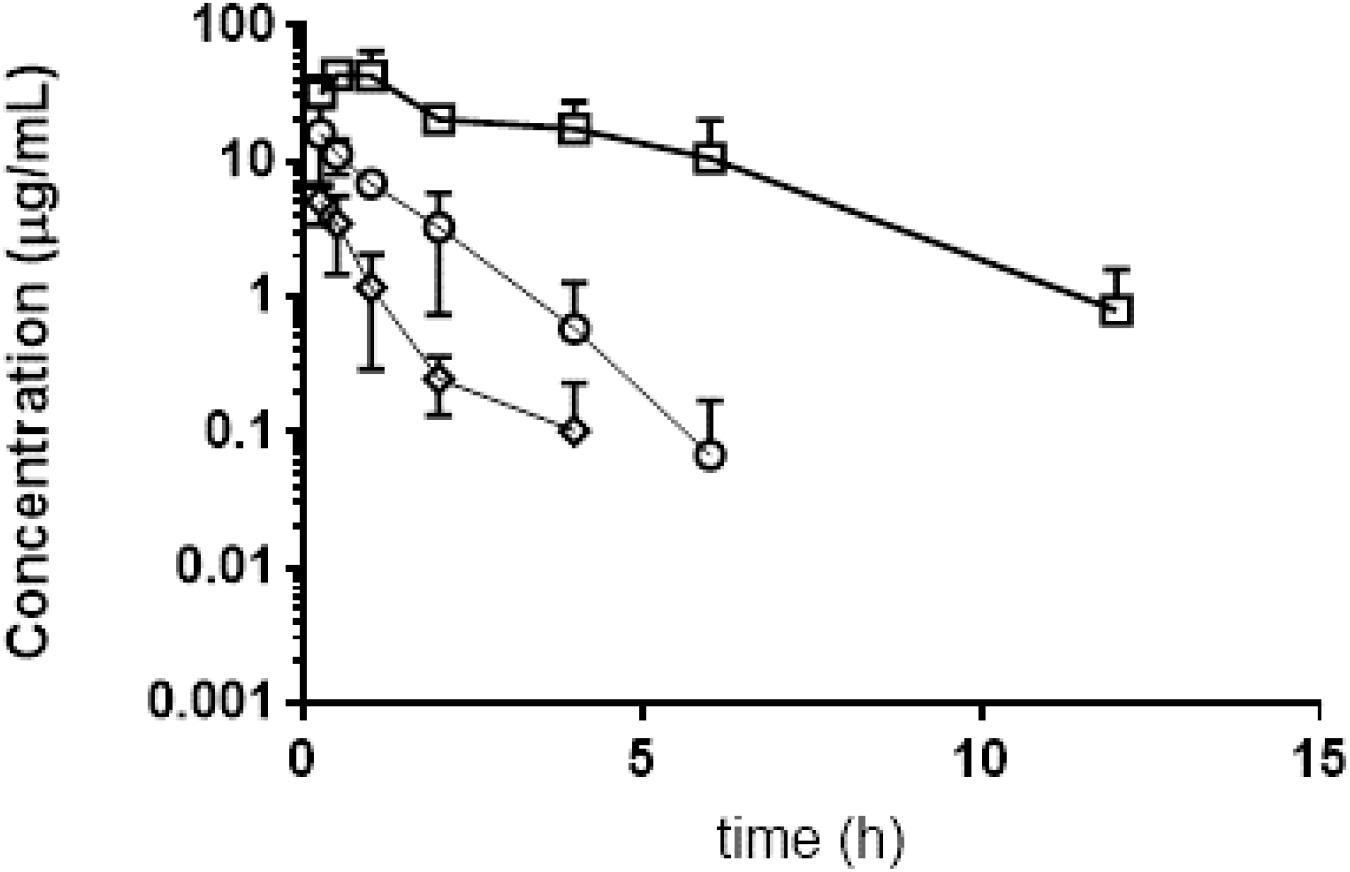
Plasma concentration of TIZ following administration of NTZ to hamsters at doses of 485mg/kg (⍰), 98.1mg/kg (⍰) and 25.5mg/kg. Three animals per dose group were included error bars represent the standard deviation.

**Supplementary Fig. 4:**
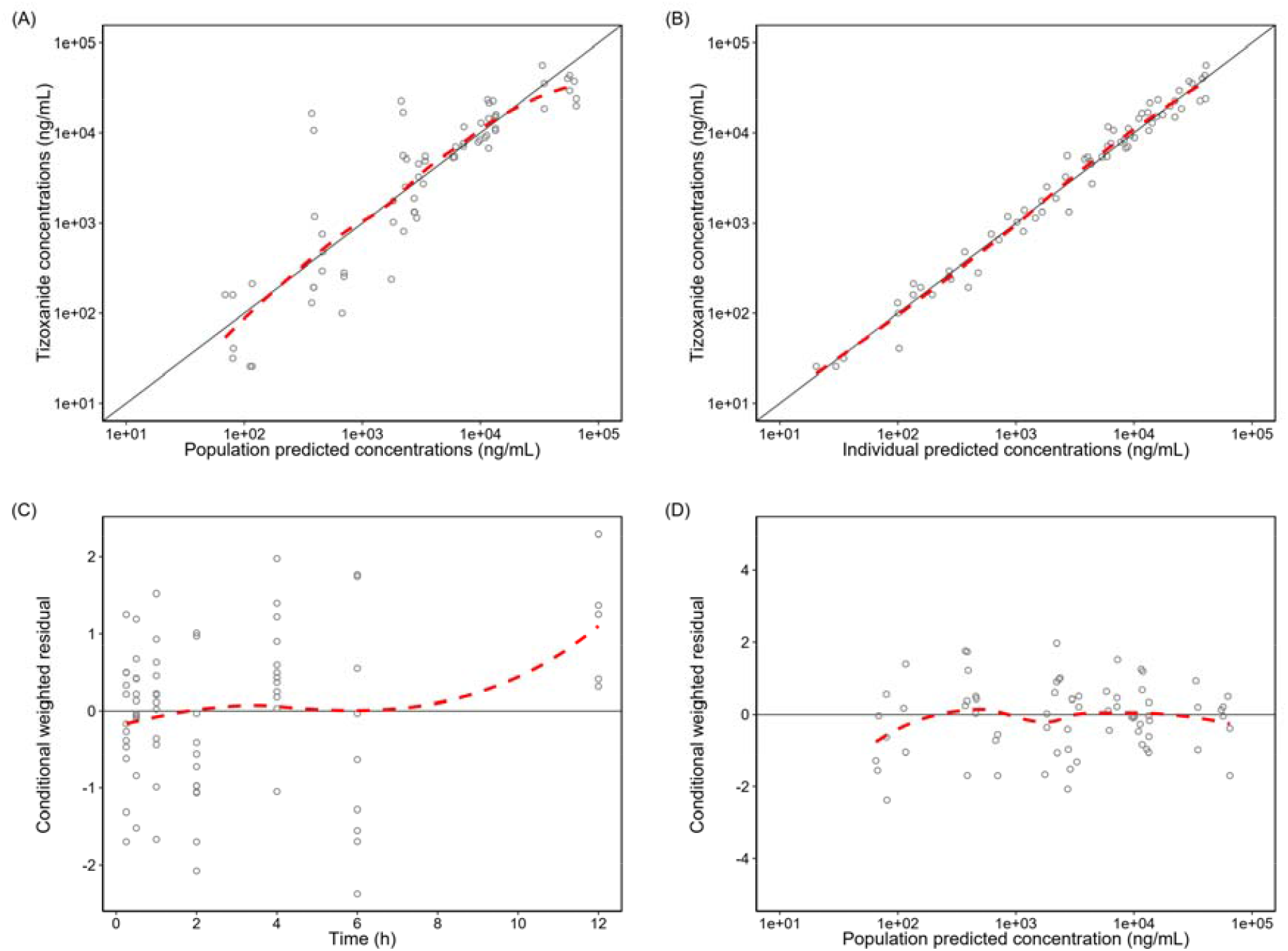
Goodness-of-fit diagnostics of final nitazoxanide population pharmacokinetic model in hamster. (A) Observed tizoxanide concentrations vs population predictions, (B) observed tizoxanide concentrations vs individually predicted concentrations, (C) conditionally weighted residual vs time, and (D) conditionally weighted residual vs population predictions. The open circles represent the observed tizoxanide concentrations. The solid black lines represent the line of identity and the dashed red lines represent a local polynomial regression fitting of all observations (i.e. trend line).

**Supplementary Table 1:** Plasma lung and nasal turbinates concentrations of TIZ after administration of multiple dose of NTZ. PK realized after 3 days of nitazoxanide administered three times a day, at the end of the dosing interval (trough concentrations). Data represent individual values (Details in Supplementary Data 4). Symbols $ and £ represent respectively 4 or 5 values below the limit of quantification.

|  | Time post-treatment | Plasma (µg/mL) | Lung (µg/g) | L/p ratio | Nasal turbinates (µg/mL) |
| --- | --- | --- | --- | --- | --- |
| Multiple Dose : 2.8mg/kg/day TID (at 3 dpi) | 12 hours | 0.01£<br>(0.03µM) | 0.10 ; 0.11 \$<br>(0.39 ; 0.41µM/g) | 1279.5 | 0.01£<br>(0.06µM) |

**Supplementary Table 2:** Population pharmacokinetic parameters from the final model of NTZ in hamster.

| Parameter | Population estimates <sup>a</sup><br>(%RSE) <sup>b</sup> | 95%CI<sup>b</sup> | IIV <sup>a</sup> [%CV]<br>(%RSE) <sup>b</sup> | 95%CI <sup>b</sup> |
| --- | --- | --- | --- | --- |
| F | 1 (fixed) | - | - | - |
| CL/F (L/h) | 0.651 (11.9) | 0.504-0.866 | 40.9 (26.5) | 14.8-53.5 |
| V/F (L) | 0.128 (48.3) | 0.044-0.359 | - | - |
| Q/F (L/h) | 0.391 (73.5) | 0.134-1.373 | 55.7 (156) | 26.8-220 |
| VP/F (L) | 0.262 (32.5) | 0.120-0.426 | - | - |
| k<sub>a</sub> (h<sup>-1</sup>) | 1.74 (56.2) | 0.941-5.945 | 99.6 (33.4) | 40.8-267 |
| σ | 0.151 | 0.088-0.204 | - | - |

<sup>a</sup> Population mean values, inter-individual variability (IIV) were estimated by NONMEM. The coefficient of
variation (%CV) for IIV were calculated as $100 \times \sqrt{\exp(\text{estimate}) - 1}$ .
<sup>b</sup> Relative standard error (%RSE) was calculated as $100 \times \left( \frac{SD}{\text{Mean value}} \right)$ from the non-parametric bootstrap results (n=1,000).
The 95% confidence interval (95%CI) is presented as the 2.5 to 97.5 percentiles of bootstrap estimates.
(F : bioavailability ; CL/F : oral clearance ; V/F : volume of distribution of the central compartment ; Q/F : intercompartmental clearance ; VP/F : volume of distribution of peripheral compartment ; K<sub>a</sub> : absorption rate constant ; σ : residual variability)

**supplementary Table 3:**
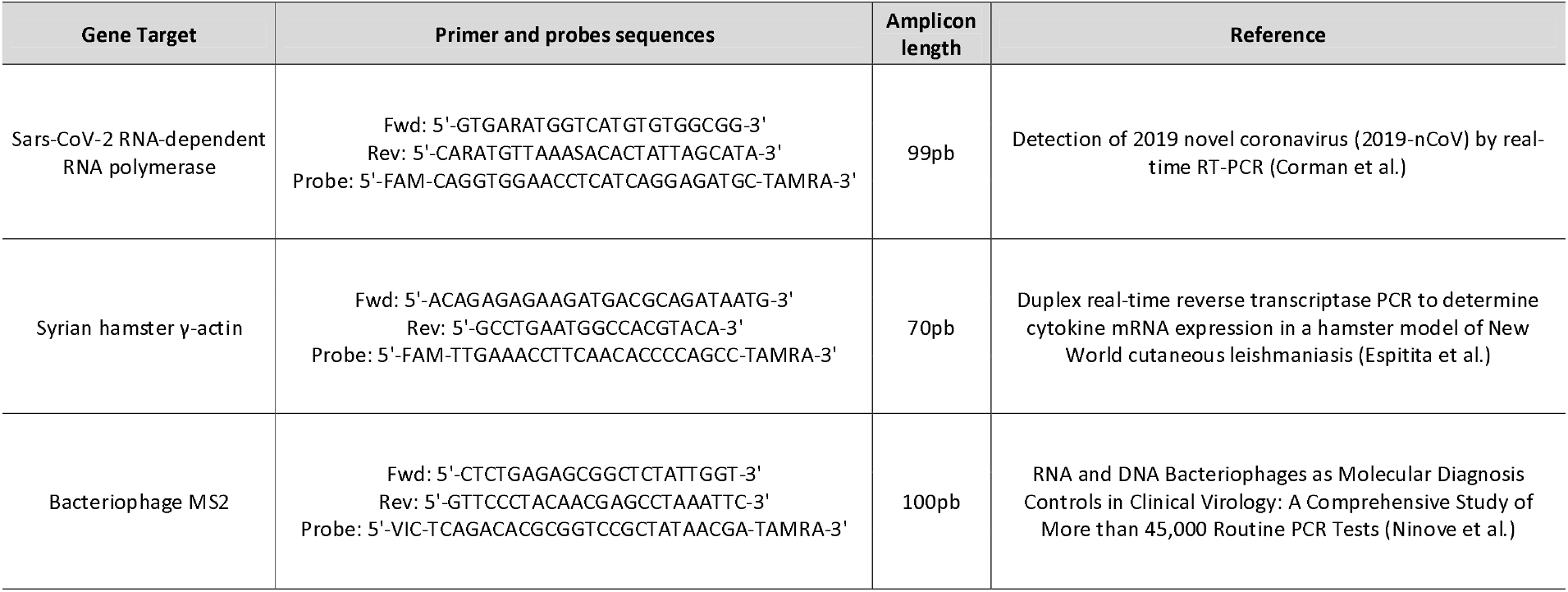
(RT)-qPCR systems.

**supplementary Table 4:** Histopathological semi-quantitative lung inflammation scoring system

| Lesion | Description | Intensity | Score |
| --- | --- | --- | --- |
| Interstitial pneumonia | 1 or 2 foci with 10-20 cells or small area with two-fold thickening of alveolar septa | Mild | 1 |
|  | 3 to 5 foci with 10-30 cells or widespread areas with two-fold thickening of alveolar septa | Moderate | 2 |
|  | 5 foci of 10-50 cells or widespread areas with two-fold or three-fold thickening of alveolar septa throughout the lung | Marked | 3 |
|  | 5 foci of 10-100 cells or widespread areas with three to fourfold-thickened alveolar septa throughout the lung | Severe | 4 |
| Bronchitis | 1 or 2 bronchi section(s) filled with rare necrotic/inflammatory cells or partially surrounded by scarce inflammatory cells | Mild | 1 |
|  | 3 to 5 bronchi filled with necrotic/inflammatory cells or partially surrounded by a few inflammatory cells | Moderate | 2 |
|  | 6 to 10 bronchi filled with necrotic/inflammatory cells, partially or sub-completely surrounded by inflammatory cells | Marked | 3 |
|  | Numerous bronchi filled with inflammatory or cellular debris or completely surrounded by numerous inflammatory cells | Severe | 4 |
| Endothelitis, vasculitis | Absent | - | 0 |
|  | Present | - | 1 |
| Hemorrhagic necrosis | Absent | - | 0 |
|  | Focal to multifocal | Mild to moderate | 1 |
|  | Coalescing to extensive necrosis | Severe | 2 |

**Supplementary Table 5:**
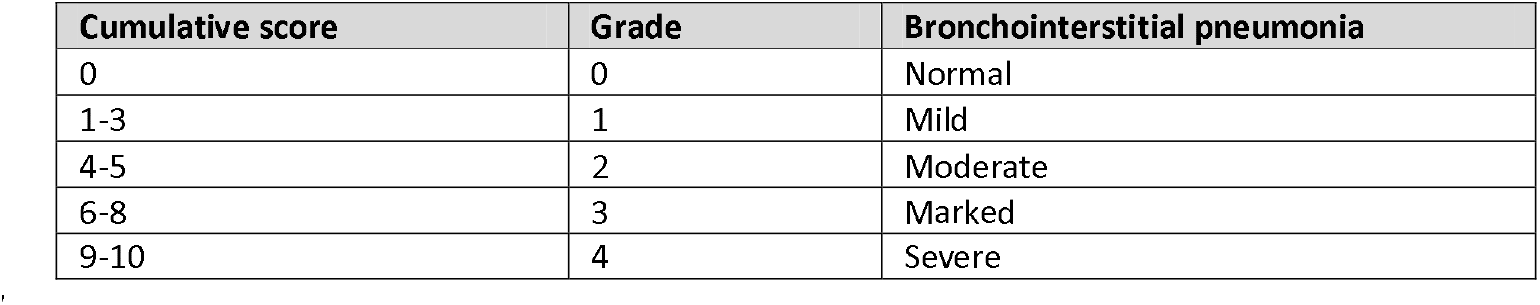
Histopathological lung inflammation semi-quantitative grading

